# Two molecules of Has1 RNA helicase function simultaneously in the biogenesis of small and large ribosomal subunits

**DOI:** 10.1101/594762

**Authors:** Sivakumar Vadivel Gnanasundram, Isabelle C. Kos-Braun, Martin Koš

## Abstract

The RNA helicase Has1 is involved in the biogenesis of both small and large ribosomal subunits. How it performs these separate roles is not fully understood. Here we provide evidence that two molecules of Has1 are recruited and temporarily present in the same time in 90S pre-ribosomes. We identified multiple Has1 binding sites in the 18S, 5.8S and 25S rRNAs. We show that while the Has1 catalytic activity is not required for binding to 5.8S/25S region in pre-rRNA, it is essential for binding to 18S sites. After the cleavage of pre-rRNA at the site A2 Has1 remains associated not only with pre-60S but unexpectedly also with the pre-40S ribosomes. The recruitment to 90S/pre-40S and pre-60S ribosomes is mutually independent. Our data reconcile some seemingly contradictory observations about Has1 function in ribosome biogenesis.

## INTRODUCTION

Ribosome biogenesis is a complex, highly dynamic major cellular metabolic process in all living cells. In the yeast Saccharomyces cerevisiae this pathway begins with the transcription of a large ribosomal RNA (rRNA) precursor, the 35S pre-rRNA in the nucleolus by RNA polymerase I. This large pre-rRNA is further processed into mature 18S, 5.8S and 25S rRNAs through a complex series of endo-and exonucleolytic cleavages and base modifications (methylations and pseudouridylations). The final maturation process takes place in the cytoplasm, where mature ribosomes catalyze the translation of mRNA into proteins. Over 200 non-ribosomal proteins, ~80 ribosomal proteins and at least 70 small nucleolar RNAs (snoRNAs) are involved in this dynamic process (1–7)

Among the accessory ribosome biogenesis factors are 19 RNA helicases, a large group of highly conserved enzymes possessing the capability to catalyze the unwinding of double-stranded RNA (dsRNA) by utilizing the energy derived from the binding and hydrolysis of ATP. These molecules are ubiquitously found in all kingdoms of life and participate in most of the processes of RNA metabolism (8, 9). Plausible functions of RNA helicases in ribosome biogenesis includes unwinding of snoRNA-pre-rRNA base pairing, remodelling of protein-RNA interactions, pre-rRNA folding and structural rearrangements. The *S. cerevisiae* Has1 is an essential protein belonging to the DEAD box family of RNA helicases. It is one of three factors (Has1, Rrp5 and Prp43) known to participate in the maturation of both the subunits. Has1 was implicated in the biogenesis of both 40S and 60S subunits, as its depletion led to the loss of 20S pre-rRNA with accumulation of 35S pre-rRNA and aberrant 23S pre-rRNA, as well as a delay in the processing of 27SB pre-rRNA (10). The ATP dependent unwinding activity of Has1 is essential for its function *in vivo* (11). Additionally, Has1 depletion led to the accumulation of snoRNPs (including U3 and U14, snR10 and snR63 snoRNAs) associated with 90S/60S pre-ribosomal particles suggesting that Has1 is required for the release of some snoRNAs (12). Affinity purifications and proteomic analysis of pre-ribosomal particles indicated that Has1 is associated with the 90S and several pre-60S particles (13–17).

To date only the role of Has1 in the 60S ribosome biogenesis has been extensively studied (18). This report suggests that the presence of Has1 in early pre-60S particle is dependent on the L7, L8 and the set of A3 factors. The authors showed that Has1 binding to pre-60S could occur in an ATP-independent manner and trigger the exonucleolytic trimming of 27S A3 pre-rRNA to generate the 5’ end of 5.8S rRNA. It also suggests that the enzymatic activity of Has1 is needed for the efficient assembly of r-proteins L26, L35 and L37 as well as for the cleavage of 27SB pre-rRNA. However, Has1 role in the 90S and biogenesis of 40S ribosomal subunit remains largely unexplored.

Here we analyzed in detail the protein composition of pre-ribosome purified directly via Has1 and show that it is present not only in 90S, pre-60S and but also pre-40S particles. Using RNA-protein crosslinking, we identified multiple binding sites of Has1 on 18S, 5.8S and 25S rRNAs. Our data corroborate well the RNA binding sites reported in a new study published during preparation of this manuscript (19). Furthermore, we found that two copies of Has1 are temporarily present in 90S pre-ribosomes and remain associated with both pre-40S and pre-60S after the cleavage of pre-rRNA at the site A2.

## RESULTS

### Has1 is a component of distinct pre-40S and pre-60S particles

Has1 RNA helicase was implicated in the biogenesis of 40S and 60S subunits. It was identified in the purified 90S and early pre-60S pre-ribosomes (10, 11, 13, 16–18). However, detailed protein composition of pre-ribosomes purified directly via Has1 has not been reported. To understand whether the presence of Has1 in both 90S and pre-60S particles is due to separate and independent roles in the ribosome biogenesis process, we reanalyzed in detail the pre-ribosomal complexes co-purifying with Has1 using mass spectrometry and RNA analysis. We created a yeast strain expressing a Has1-FTP with the C-terminal affinity tag consisting of the FLAG tag, TEV cleavage site and a protein-A tag. The Has1-FTP purified pre-ribosomes consisting of a mixture of early 90S, pre-40S and predominantly pre-60S factors as well as ribosomal proteins (Figure 1A). Has1 was most tightly associated with pre-60S ribosomes, as washing the affinity purified pre-ribosomes with a series of salt washes of increasing stringency, NaCl [100 mM-1000 mM] or MgCl2 [5 mM-100 mM] led to dissociation of the pre-90S and pre-40S factors but not pre60S factors (Supplementary Figure 1 and data not shown). In total we identified 147 proteins that reproducibly co-purified with Has1-FTP in three independent experiments. Their relative abundance to the bait (iBAQ values) is shown in the Figure 1B and Supplementary Table 1. The complex mixture of ribosome biogenesis factors from different stages indicated that Has1 was likely present in at least three different particles, broadly corresponding to 90S, pre-60S and pre-40S pre-ribosomes. This conclusion was further corroborated by the analysis of pre-rRNAs co-purifying with the Has1 that showed that Has1 strongly purified the 35S, 27SA/B and 7S, and 20S pre-rRNAs, which are components of 90S, pre-60S and pre-40S pre-ribosomes respectively (Figure 1C left).

**Figure 1.**
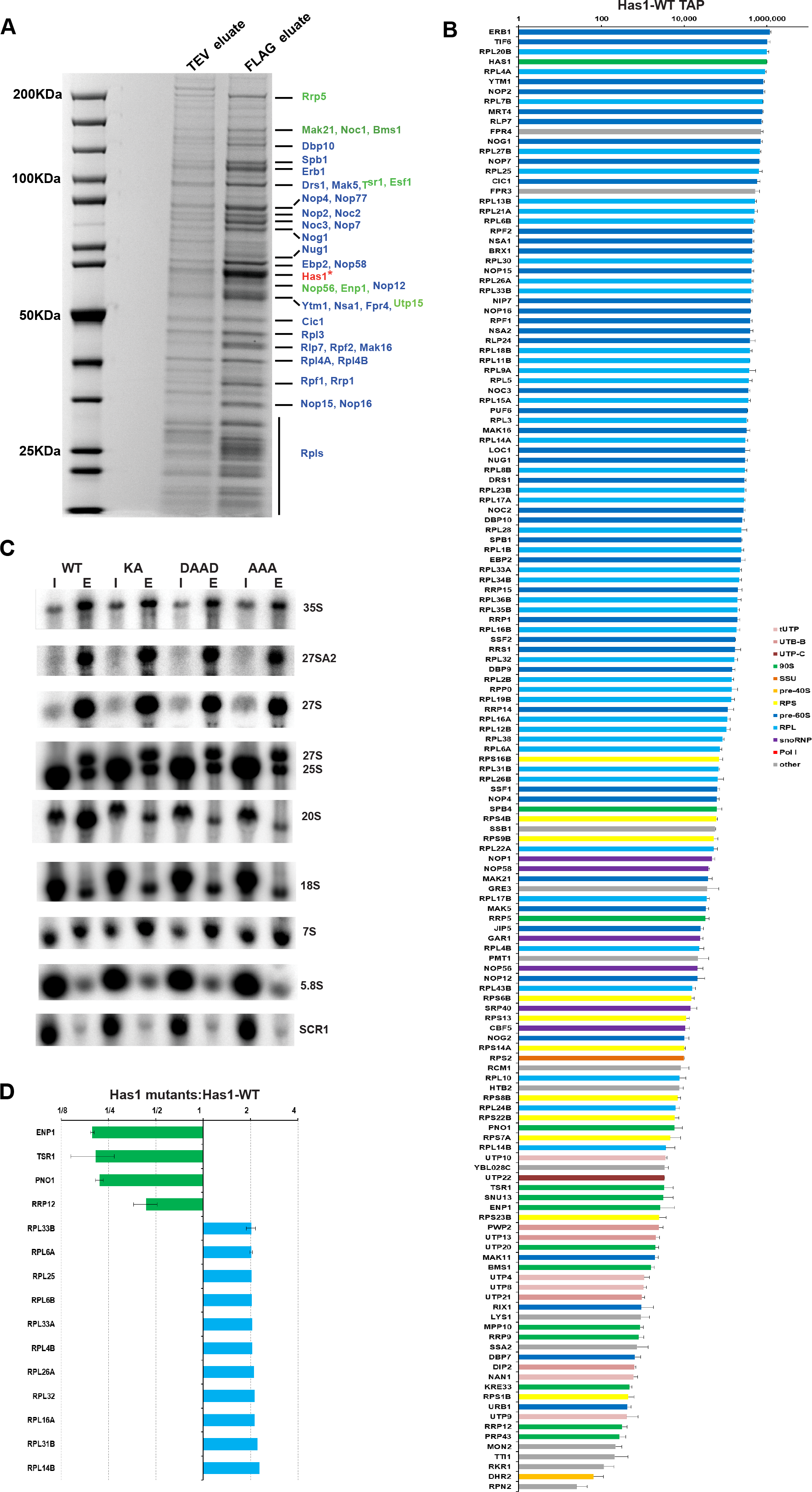
Has1 is a component of both small and large pre-ribosomal particles. SDS PAGE analysis of the proteins co-purified with the wild type Has1-FTP. The TEV (10%) and FLAG eluates were resolved in 4-12% gradient SDS-PAGE and stained by Colloidal Coomassie Blue. The bait protein Has1 is marked with the asterisk. The marked proteins were identified by MALDI-TOF mass spectrometry. In blue are pre-60S factors and in green 90s/pre40s factors. B) Protein composition of the wild type Has1 pre-ribosomes. The iBAQ values normalized to bait are plotted (Has1 value was fixed to 10^6^). C) Northern blotting analysis of RNA composition of pre-ribosomes affinity purified via Has1-FTP. D) Changes in protein composition of the pre-ribosomes purified via the catalytically inactive Has1 mutants. The SILAC H/L ratios normalized to bait are shown.

Dembowski *et al.* found that also the catalytically inactive Has1 could be purified with the Rpf2 and Rrp5 containing pre-ribosomes (18), however, in which pre-ribosomal stages can catalytically inactive Has1 enter was not analyzed in detail. Here we analyzed directly the composition of the pre-ribosomes purified via the catalytically inactive Has1 mutants using SILAC mass spectrometry. We created a yeast strain with the endogenous *HAS1* gene under the control of repressible TetO7 promoter (TetO7-Ubi-Leu-3HA Has1), allowing fast depletion of the wild type Has1 upon addition of doxycycline to culture (20). This strain was transformed with plasmids carrying either the wild type or catalytic mutants of Has1 fused at their C-terminus to the FTP affinity tag under the endogenous *HAS1* promoter. Mutations were made in the conserved RNA helicase motives I (mutation K92A [KA]), II (E197A [DAAD]) and III (S228A/T230A [AAA]) which are critical for the ATP binding, ATP hydrolysis and helicase activity respectively. We hypothesized, that the Has1 mutants defective in different steps of the RNA helicase ATP cycle might arrest the maturation of pre-ribosomes at different states. The strain expressing the plasmid born wild type Has1 was grown in a heavy isotope media and mutants expressing strains in light media. The wild type Has1 expressed from the genomic copy was depleted for 4 hours. The wild type culture was mixed with each mutant culture in equal ratios and Has1 was purified via the FTP tag and the FLAG eluate was analyzed by mass spectroscopy. The overall protein composition of the Has1 pre-ribosomes was very similar between the three mutants, indicating that the three different catalytic mutants did not arrest pre-ribosomes at distinct stages of biogenesis (Supplementary Table 2). Surprisingly, there were also only few significant changes compared to wild type Has1 pre-ribosomes (Figure 1D). Four 90S/pre-40S biogenesis factors, Enp1, Tsr1, Pno1 and Rrp12 were clearly reduced, while several ribosomal proteins of the large subunit were enriched in the mutant Has1 pre-ribosomes (Figure 1D).

Next, we analyzed the pre-rRNA composition of pre-ribosomes purified with either wild type or catalytically inactive Has1. As the Has1 catalytic activity is required for the pre-rRNA cleavage at the site A2, which is essential for formation of pre-40S particles, the potential interaction of inactive Has1 with pre-40S pre-ribosomes cannot be analyzed under depletion conditions, since no pre-40S particles are formed. We therefore purified the plasmid born wild type or mutant Has1-FTP without depletion of the endogenous wild type Has1 protein and analyzed the pre-RNA composition by northern blotting (Figure 1C right). As expected, the wild type Has1 purified well the primary transcript 35S pre-rRNA as well as intermediates of both 40S and 60S subunits the 20S and 27S pre-rRNAs. All the catalytic mutants also purified 27S pre-rRNAs equally well as the wild type protein, but association with 20S pre-rRNA containing pre-ribosomes was strongly reduced. Intriguingly, the Has1 mutants also purified 27S-A2 pre-rRNA, which can be produced only in the presence of fully active wild type Has1. This indicated that either the Has1 is turned over and can be exchanged in the pre-ribosomes (the mutant Has1 can binds to the 27S-A2 containing particle that was produced in the presence of the wild type Has1) or that two molecules of Has1 might be present in the early 90S pre-ribosomes, which turned out to be the case, as we show later. To determine whether Has1 is present in distinct particles the affinity purified wild type or Has1-DAAD mutant pre-ribosomes were fractionated by sucrose gradient centrifugation. Both proteins and RNA were extracted from each fraction and analyzed by electrophoresis (Figure 2A, 2B). Based on the pre-rRNA species present in each fraction, the wild type Has1 clearly purified the pre-40S (fraction 6) and pre-60S and 90S pre-ribosomes (fraction 8, 9 and 10) respectively. Several of the proteins in the fraction 6 were present in approximately stoichiometric amounts. Using MALDI-TOF these proteins were identified as Tsr1, Ltv1, Has1, Rio2, Pno1 (Figure 2A). Contrary to the wild type, the Has1-DAAD purified only pre-60S and 90S intermediates and no pre-40S was detectable (Figure 2B), confirming that catalytically inactive Has1 cannot be recruited to pre-40S particles. It is important to note, that small amounts of 20S pre-rRNA were also present in the fraction 9 in addition to 35S and 27S pre-rRNA. This presumably represents a transient state of 90S pre-ribosomes immediately after the cleavage at A2 site when both 20S and 27S-A2 pre-rRNAs are still present in one particle.

**Figure 2.**
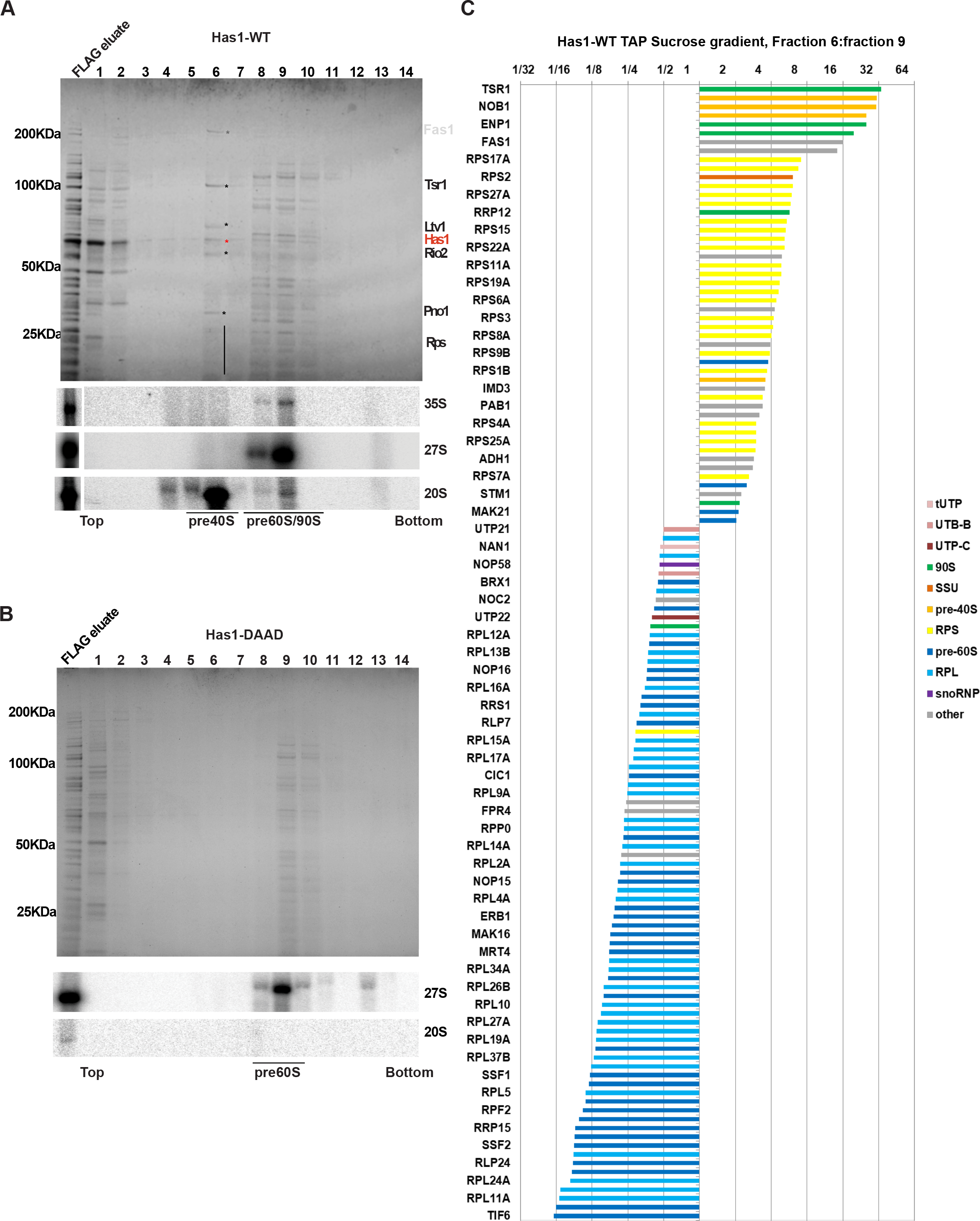
Has1 is present in distinct pre-ribosomes. A) Top: Affinity purified Has1 wild type material was fractionated on sucrose gradient. and resolved by SDS-PAGE and Coomassie stained. Bottom: northern blotting analysis of RNA in each fraction. B) Same as in A but Has1-DAAD mutant. C) Spike-in SILAC analysis of protein composition of fractions 6 and 9 from the purification via the catalytically inactive Has1-DAAD.

To analyze the protein composition of different fractions more comprehensively, we used spiked-in SILAC approach. Briefly, the gradient fractions (6 and 9) of Has1-FTP purified from a yeast culture grown in the light isotope media were spiked with equal amounts of the FLAG eluate (complete, not separated on a gradient) from a culture grown in a heavy isotope media. This allowed us to directly compare the fractions and identify which proteins are enriched or lost in each fraction compared to total FLAG eluate (input). In agreement with the pre-rRNAs present in each fraction, the lighter fraction 6 consists of predominantly pre-40S factors and small ribosomal proteins whereas the pre-60S and 90S factors and large ribosomal proteins are predominant in the slower sedimenting fraction 9 (Figure 2C and Supplementary figure 2). We conclude that the wild type Has1 is present in all pre-90S, pre-40S and pre-60S pre-ribosomes, while the catalytically inactive Has1 can associate with pre-90S and pre-60S, but not pre-40S pre-ribosomes.

### Has1 binds to both 18S and 25S regions in the pre-rRNA

As an RNA helicase, it is likely that Has1 contacts RNA directly during its function in ribosome biogenesis. To identify the Has1 binding sites on the pre-rRNA, we performed the UV cross-linking and analysis of cDNA (CRAC) (21) using strains expressing either the wild type Has1 or the catalytically inactive Has1-DAAD mutant. Both proteins crosslinked to multiple regions in the pre-rRNAs (Figure 3A). The wild type Has1 crosslinked to 3’ major (3’M) domain of 18S rRNA (helices 31-41), 5.8S rRNA and ITS2 region, and to 5’end of 25S rRNA (helices 16, 17 and 21, 22). The Has1-DAAD crosslinking produced overall ~3 times fewer reads compared to wild type. The crosslinking pattern remained similar to wild type at the 5’ end of 25S rRNA (helices 17-22), but was strongly reduced in the 5.8S and 18S region. Several new peaks were present in the Has1-DAAD crosslinking profile (Figure 3B and 3C). Additional peak in the 18S region, representing crosslinking to the helices 17 and 18. Another two regions in the 25S rRNA, corresponding to helices 79 and 89 were strongly amplified, however, these are known contaminating peaks from previous studies (22, 23) and were probably over-amplified due to overall reduced crosslinking efficiency of the mutant Has1-DAAD. In order to obtain better sensitivity of crosslinking, we repeated the crosslinking experiment using 4-thiouridine (4-TU) in the culture media. The wild type Has1 produced very strong crosslink (about 9x higher number of reads) with the profile very similar to the standard CRAC experiment (Figure 3A). Interestingly, a new crosslinking site was detected in the 18S rRNA region, helices 4 and 6. This crosslinking pattern is in very good agreement with the recently published Has1 PAR-CRAC (19). Unfortunately, we were not able to crosslink the mutant Has1-DAAD using 4-TU for to us unknown reasons. The binding sites of Has1 in 18S rRNA are in close proximity to the known binding sites of other pre-40S processing factors (Enp1, Ltv1, Nob1 and Rio2), while the binding sites of Has1 in 25S rRNA are in the vicinity of binding sites of Erb1, Nop7, Nop12 and Nop15 (22, 23) (Supplementary table 3 and 4). These results support the observation that Has1 can participate in the rRNA processing by directly interacting with rRNA of both small and large ribosomal subunits.

**Figure 3.**
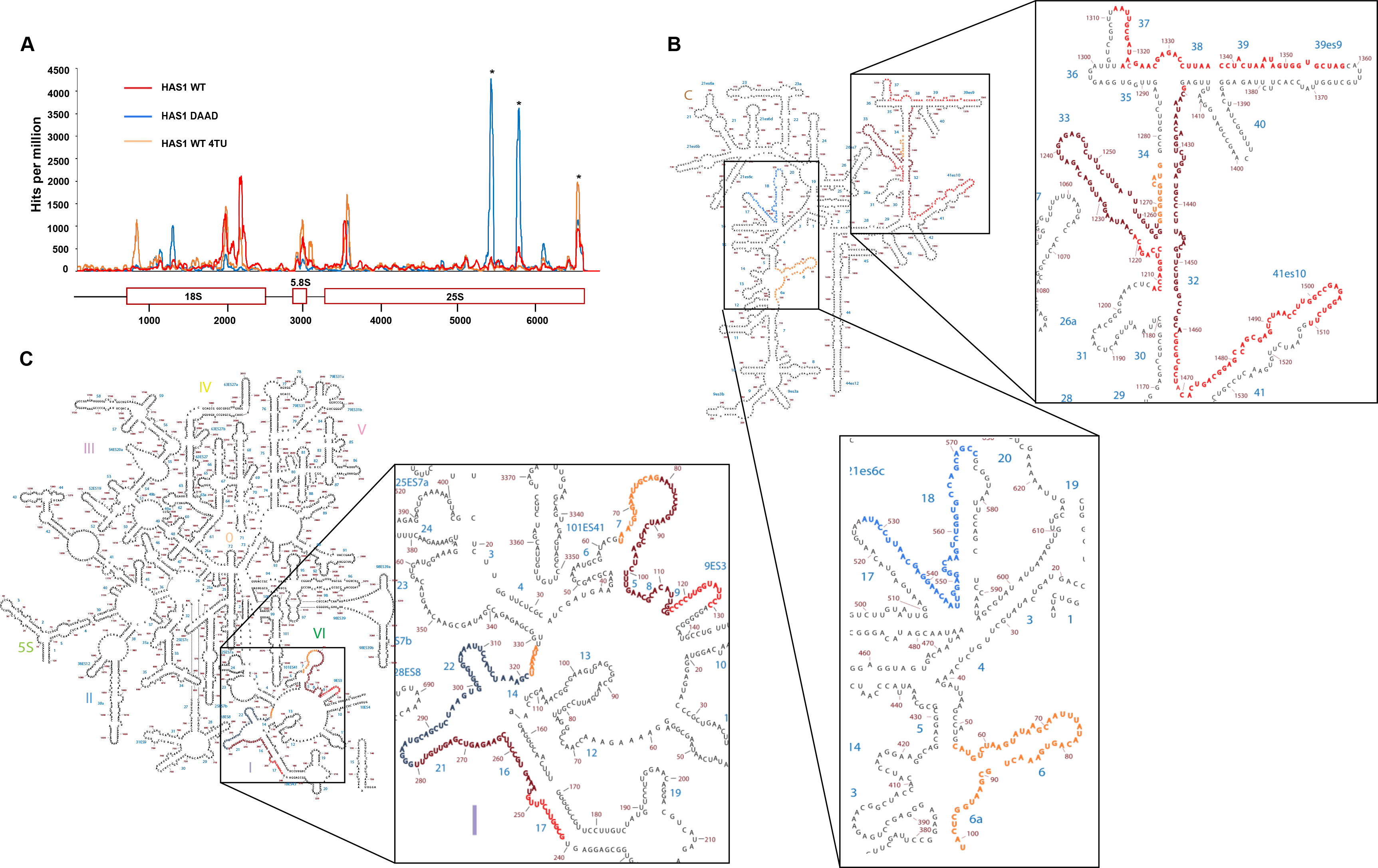
Has1 crosslinks to multiple regions of pre-rRNAs. A) Plot showing the distribution of reads coverage across pre-rRNA. Average number of hits from two independent experiments was plotted. Reads per 1 million total reads are shown. Asterisks indicate common contaminating peaks. B) and C) Binding sites of both Has1 WT and DAAD mutant were mapped on the secondary structure of 18S (B) or 5.8S and 25S rRNAs (C): red-Has1 wild type, orange-Has1 wild type 4TU, blue-Has1 DAAD. The overlapping sites of Has1 wild type from standard and 4TU CRAC are shown in dark red. The overlapping sites of Has1 wild type and Has1-DAAD are in dark blue.

### Two Has1 molecules temporarily coexist in 90S complex

Since Has1 co-purified early 90S, pre-40S and pre-60S factors and also interacted with multiple sites in both 18S and 25S rRNA, it is plausible that Has1 is present in two independent copies in the pre-ribosomes. This would also explain the observed purification of 27S-A2 by Has1-mutants (Figure 1C). To investigate this hypothesis, a yeast strain was created with the endogenous Has1 tagged with 3HA and expressing a plasmid born Has1-FTP. In principle if two copies of Has1 are present in the same pre-ribosome particle, then the affinity purification through one copy of Has1 should co-purify the other. We purified pre-ribosomes via the Has1-FTP and analyzed them for presence of the Has1-3HA by Western blotting. The tRNA binding protein Arc1 was used as the negative control. As can be seen in the Figure 4A, the 3HA-Has1 clearly co-purified with the Has1-FTP, confirming that two copies of Has1 RNA helicase are simultaneously present in the same pre-ribosomes (Figure 4A and 4B). The same result was obtained also in a strain co-expressing a Has1-GFP and Has1-FTP (Supplementary Figure 3).

**Figure 4.**
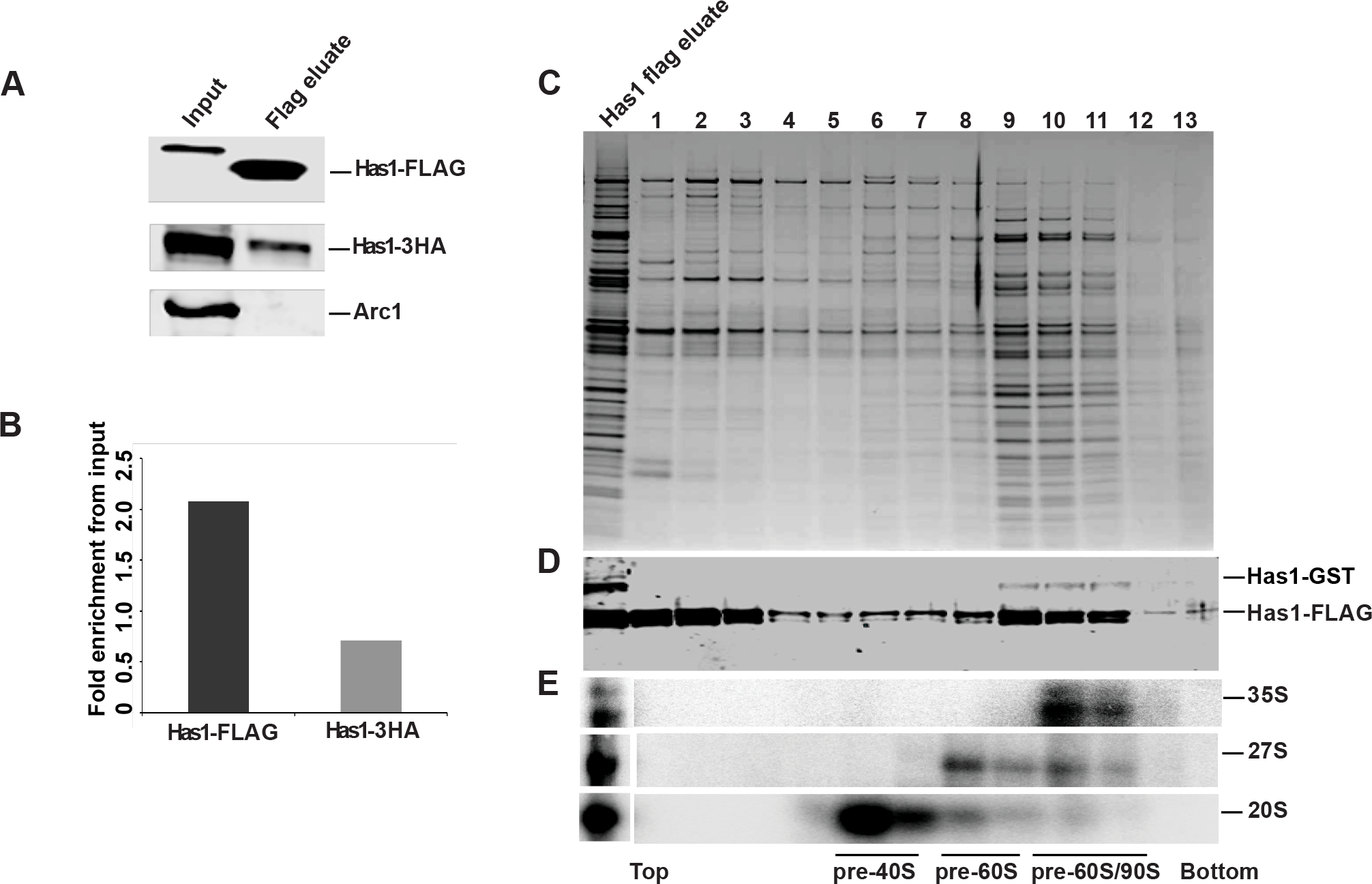
Has1 is present in two copies in 90S pre-ribosomes. A) Western blot analysis of Has1-FTP purification form the yeast strain expressing both 3HA-Has1 and Has1-FTP using anti-3HA, anti-FLAG and anti-Arc1 antibodies as indicated. The change in the molecular size of Has1-FLAG is due to the TEV cleavage during tandem affinity purification. Chart showing the average fold enrichment of Has1-3HA, Has1-FLAG based on the quantification of data from two independent experiments. C) Sucrose density gradient analysis of the Has1-FTP purified material from a strain expressing both Has1-GST and Has-FTP. Proteins isolated from each fraction were resolved by SDS PAGE and detected by v Coomassie staining. D) Western analysis of the fractions using the anti-Has1 and anti-GST antibodies. E) Northern analysis of RNA in each fraction. The fractions corresponding to different pre-ribosomes based on the pre-rRNA content are indicated below the image.

To determine, which pre-ribosomes contain the two copies of Has1 we fractionated pre-ribosomes purified via the Has1-FTP by size using the sucrose-density gradient centrifugation (Figure 4C, 4D). To be able to better distinguish different Has1 copies in the Western analysis, we used a yeast strain with the genomic copy of Has1 tagged with GST at the C-terminus and expressing the plasmid borne Has1-FTP. Corroborating the previous observation, Has1-GST was co-purified with the Has1-FTP (Figure 4D). Importantly, the Has1-GST was detected only in the fractions 10-11 corresponding to the 90S pre-ribosomes (based on the presence of 35S pre-rRNA) and in the fraction 9 corresponding to the pre-60S particles contains 27S pre-rRNA (Figure 4E). Notably, the fractions 9-10 contained in addition to 27S pre-rRNA also small amounts of 20S pre-rRNA (Fig 4E top) suggesting that the two copies of Has1 remain temporarily in the same pre-ribosomes after the A2 cleavage.

### Recruitment of Has1 to pre-40S and pre-60S/90S is mutually independent

To understand whether Has1 recruitment into pre-ribosomes occurs at one or more time points, we employed a set of rDNA truncations plasmids. Briefly, yeast strains expressing Has1-FTP were transformed with a set of multicopy plasmids that carry a copy of the whole rDNA unit (with the native Pol I promoter) truncated at different distance from the transcription start site by introduction of a hybridization tag (corresponding to a MS2 binding site) followed by the native endogenous termination region from the 3’ETS (Figure 5A). Consequently, these truncations are expressed from the native RNA polymerase I promoter and the transcription terminates at the native terminator, ensuring the processing of the pre-rRNAs synthesized from these rDNA truncations is as physiological as possible. The rRNA truncations that co-purified with Has1-FTP were detected by northern blotting analysis using a probe complementary to the hybridization tag (Figure 5A, 5B). The SCR1 (small cytoplasmic RNA) RNA, was used as the loading control. Has1 did not associate with the rRNA truncation which contains only the 5’ ETS and showed clear enrichment only for the truncations containing the 5’ETS and 18S rRNA/ITS1 regions. However, Has1 very strongly co-purified the rRNA truncations containing the 5’ end of the 25S rRNA (1-421nt) (Figure 5C). In summary, these results suggest that Has1 is recruited to pre-ribosomes after the complete 18S rRNA is transcribed, and mainly upon transcription of the 5’end of the 25S rRNA.

**Figure 5.**
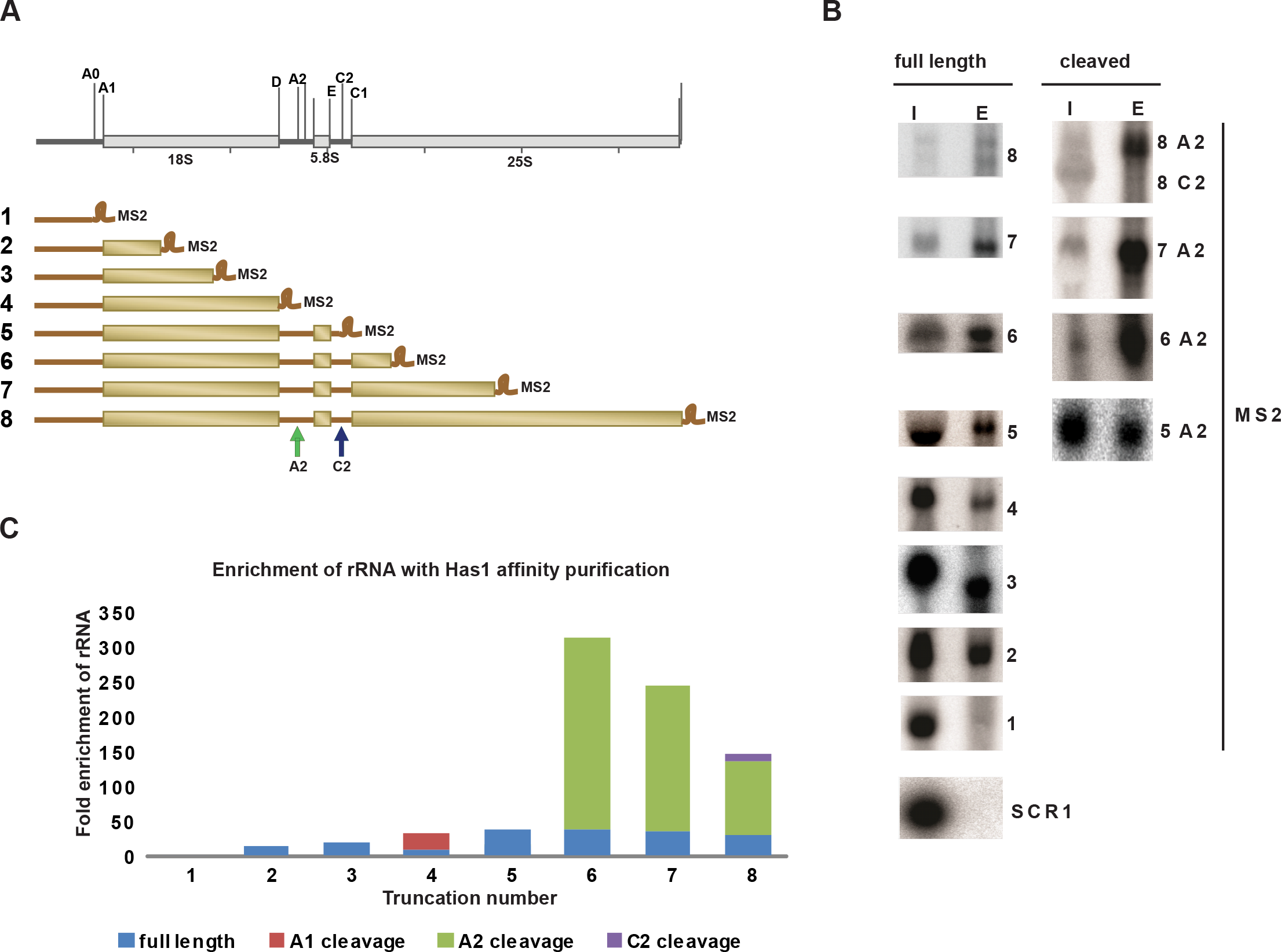
Timing of Has1 joining in the pre-rRNA processing. A) Schematic representation of the 35S rDNA and various rDNA truncations with MS2 tag used. The cleavage sites A2 and C2 are marked on the full-length truncation. B) Northern blot analysis of rRNAs purifying with Has1. Lane I was loaded with 1% of input RNA, lane E is the RNA purified with Has1. For clarity, the full length and cleaved A2 or C2 cleaved pre-rRNAs are shown in separately. C) Quantification of the fold enrichment over input of pre-rRNAs purified through Has1. The full length rRNA and the cleaved ones are indicated with different color codes.

To examine if the recruitment of Has1 to the pre-60S and pre-40S is independent, we used strains expressing truncations with the 5’end (1-421nt) of 25S rRNA together with or without the 18S rRNA, ITS1 or 5.8S regions and vice versa for the 18S rRNA (Figure 6). To allow us to detect the 5’ part of the pre-rRNA after the A2 cleavage, an internal oligonucleotide tag was inserted into the 18S rRNA sequence as previously described (24). Has1 was recruited to the 25S rRNA truncations lacking the 18S rRNA and/or ITS1 regions, however it failed to associate with truncations lacking 5.8S rRNA and intact ITS2 regions (Figure 6A and 6B). Therefore, the recruitment of Has1 to pre-60S particles does not require presence of the binding sites in the 18S rRNA. Identical results have been previously obtained by Chen et al 2017 (25). Similarly, for the binding to 18S portion of pre-rRNA, as can be seen from the Figure 6C and 6D, Has1 can bind truncations containing solely the 18S rRNA, however, the interaction is enhanced by when the ITS1 region is included.

**Figure 6.**
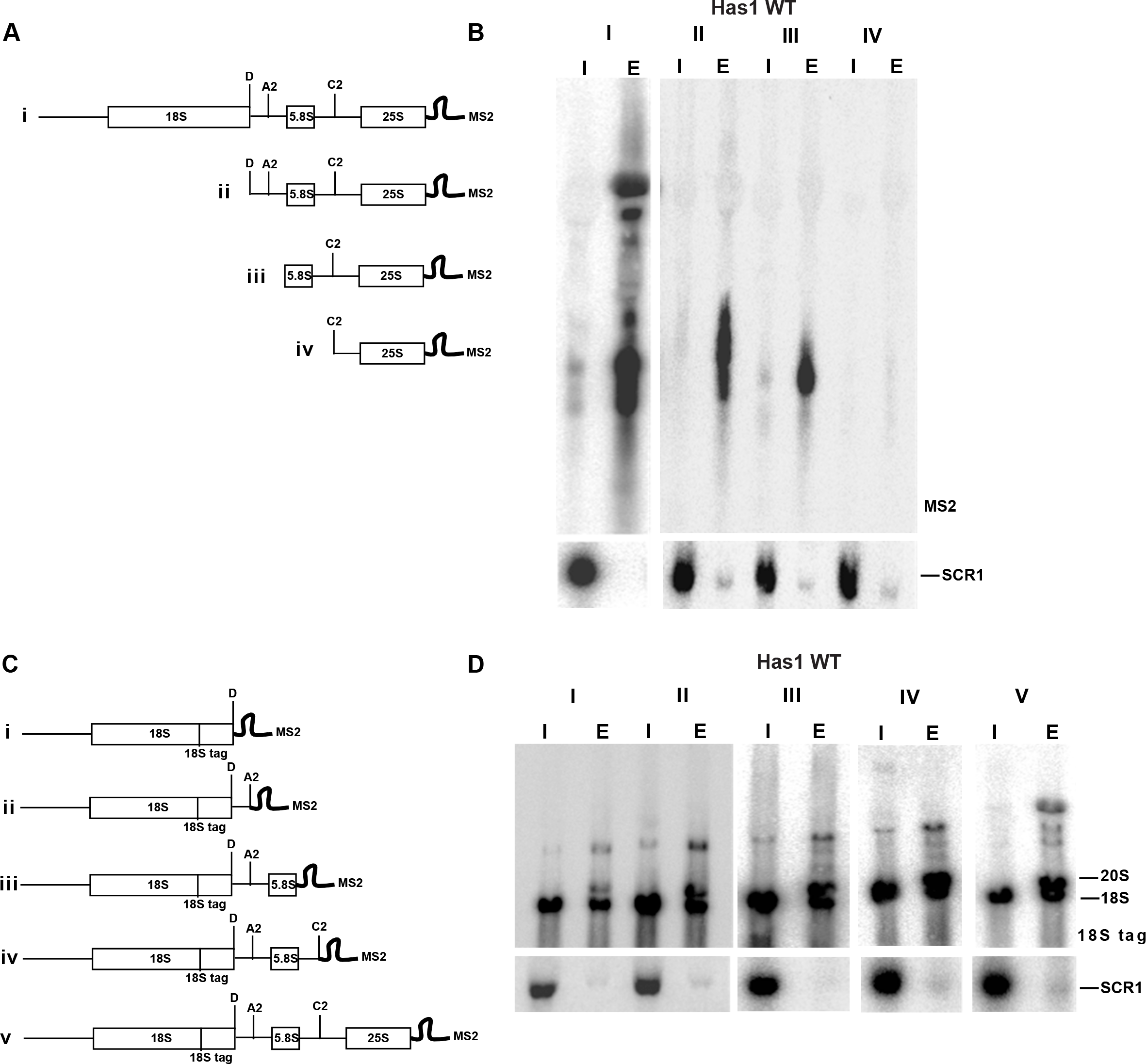
Has1 can be recruited to 18S and 25S regions in pre-rRNA independently. A), C) Schematic representation of rDNA truncations used. B), D) RNAs isolated after one-step purification of Has1 were analyzed by northern blotting using probe against the MS2 tag or 18S-tag respectively. Probing for SCR1 RNA was used as the loading control. Lanes I −1% of the input, lanes E – eluted RNA purified with Has1 FTP.

Taken together the data presented here we conclude that two molecules of Has1 are recruited to pre-ribosomes and that recruitment to the pre-90S/40S and pre-60S particles can occur independently.

## DISCUSSION

Has1 was initially identified in the affinity purified 90S and early 60S pre-ribosomal particles (10, 11, 13, 16, 17) and it was reported that depletion of Has1 led to defects in the synthesis and processing of both 18S and 25S rRNAs (10). In this study, we analyzed for the first time the composition of pre-ribosomes purified directly via affinity tagged Has1. In agreement with the previously published data, we found that Has1 purifies a mixture of 90S factors, pre-40S and pre-60S factors. It is most stably associated with the pre-60S particles as evidenced by release of 90S and pre-40S factors during washes with increasing salt concentrations. On the RNA level, wild type Has1 purified 35S, 27SA2, 27SA3, 27SB, 20S and 7S pre-rRNAs, corroborating previous reports (10, 14, 18). Interestingly, while the experiments in Figure 5 show that binding to the 18S region of the pre-rRNA is weaker compared to binding to 5.8S/25S region, Has1 strongly purifies 20S pre-rRNA. It is possible that Has1 is recruited to the pre-rRNA immediately prior to A2 cleavage, mediates the cleavage and then remains temporarily associated with the freshly generated 20S pre-rRNA. The relatively long life-time of 20S pre-rRNA (26), compared to other intermediates, might also contribute to the apparent strong signal.

The CRAC analysis revealed that wild type Has1 clearly interacts with both 18S/ITS1 and 5.8S/25S regions of pre-rRNAs. The crosslinking pattern of the Has1-DAAD catalytic mutant showed strong reduction in 18S region, in agreement with its inability to bind 90S and pre-40S particles. Our data for the wild type Has1 are in a very good agreement with the recently published PAR-CRAC analysis (19). The observed crosslinking sites in the 5.8S/25S region also coincide well with the location of Has1 in several recent cryoEM structures of pre-60S ribosomes (27–29). The binding sites of the wild type Has1 in the 18S/ITS1 region are in close proximity to the binding sites of the pre-40S assembly factors Enp1, Ltv1 and Tsr1 (22). Notably, the same factors co-sedimented with Has1 in approximately stoichiometric ratio in sucrose gradients (Figure 2A). The assembly factors Enp1 and Ltv1 were reported to interact directly and form a complex with Rps3, and are implicated in the late structural reorganization of the pre-40S (30). Interestingly, Tsr1 was implicated in the late steps of processing of 20S pre-rRNA in the cytoplasm (31). However, Has1 localization is reported to be nucleolar/nuclear, therefore Has1 must be released from this pre-40S complex before its export to cytoplasm. We don’t know whether Has1 mediates other structural changes in this pre-40S complex or participates in the release of the earlier 90S factors after the A2 cleavage. To date, Has1 was not identified in any of the published cryoEM structures of 90S or pre-40S ribosomes, presumably reflecting only transient nature of its residence in these particles (32–35).

Unexpectedly, the catalytically inactive Has1 mutants was able to purify 27S-A2 pre-rRNA, which can be produced only in the presence of catalytically active Has1 (Figure 1C). The purification of Has1 catalytic mutants was performed in a strain were both mutant and wild type Has1 were present (without depletion of the wild type Has1 protein). Therefore, there are two possible models that can explain the observed 27S-A2 pre-rRNA purification by mutant Has1: a) the mutant Has1 associates with pre-60S pre-ribosomes containing 27S-A2 and replaces the wild type Has1; b) there are two molecules of Has1 present at least temporarily in the late 90S pre-ribosomes – one that is required for A2 cleavage (catalytically active) and a second one that mediates 27S-A processing, which does not require catalytic activity (as shown by (18)). Our data offer multiple evidences in support of the later model, i.e. the action of two independent Has1 molecules during biogenesis. The already above mentioned purification of 27S-A2 by catalytically inactive Has1 mutants, the presence of Has1 in both pre-60S and pre-40S particles in sucrose gradients and also Has1 binding to sites in 18S and 25S rRNAs which are too far apart could be easily explained if two molecules of Has1 were performing those functions. Furthermore, the clear co-purification of two differently tagged Has1 molecules strongly implies simultaneous presence of two different Has1 molecules in pre-ribosomes. In sucrose gradients the two different Has1 molecules co-sedimented only with the 90S/heavy pre-60S particles (fractions 9-11 in the Figure 4D). This makes sense as only the 90S pre-ribosomes contain 35S pre-rRNA allowing simultaneous binding of two different Has1 molecules to 18S and 5.8S/25S regions. In addition, the quantification of two distinct Has1 molecules co-purification showed that only about a third of Has1-3HA present in the input co-purified with the Has1-FTP which served as bait (Figure 4A, 4B). This fits surprisingly well with the observation that only 30% of pre-rRNA is processed post-transcriptionally in the growing yeast cells and thus 35S pre-rRNA containing 90S pre-ribosomes represent only a third of all nascent pre-ribosomes (26). All this data supports the model of two Has1 molecules acting independently during ribosome biogenesis.

Can Has1 function as a dimer? Has1 was observed to self-interact in the yeast two-hybrid assays (36, 37).On the other hand, Has1 identified in the cryoEM structures of pre-60S ribosomes is present as a monomer (27–29). It is possible that Has1 is recruited as dimer but then separates and each monomer functions separately in the pre-40S and pre-60S pathways. We cannot formally exclude this possibility; however, our data do not support the model in which Has1 would be a functional dimer throughout the biogenesis. In case of a dimer, differently tagged Has1 should co-purify in stoichiometric amounts and also be present in all the fractions of the sucrose gradients (e.g. in Figure 4C), none of which we have observed.

In conclusion, our findings suggest that two independent molecules of the Has1 RNA helicase function in ribosome biogenesis. Two copies of Has1 are recruited to the pre-rRNA processing, one copy (Has1-18S) binds to the 18S rRNA and the second copy (Has1-25S) binds to both 5.8S and 5’end of 25S rRNA. Has1-18S, at the expense of ATP hydrolysis, performs structural rearrangements required for cleavage at the site A2 and remains temporarily associated with the resulting 20S pre-rRNA. It is then later released. The Has1-25S assist in the rearrangements and processing of 27S-A2/A3 and is released after the C2 cleavage.

## MATERIALS AND METHODS

### Yeast strains and plasmids

All the yeast strains and constructs used in this study are described in the supplementary tables 5 and 6. Unless mentioned all strains were constructed from the parental strain YMK118 (38). Strains for disruption or C-terminal gene tagging were created by PCR based method as described earlier. Strains with depletion of essential genes were constructed using TetO7-Ubi-DAA-3xHA cassette (20). All the yeast work and gene manipulations were done by following the standard methods (39).

### RNA Isolation and northern Blotting

The total RNA and northern blotting was performed as previously described (40, 41). Briefly Total RNA from equal amounts of cells was resolved on a 1% agarose gel and transferred to a Nylon membrane using TV400-EBK Maxi Electro blotter (Severn Biotech). The blot was hybridized with the [^32^P] 5’-end labelled oligonucleotides complementary to pre-rRNAs (listed in supplementary Table 7) and exposed to phosphorimaging plates and scanned on the FLA-7000 imager (Fuji).

### Tandem affinity purification

Tandem affinity purifications were performed as described (42). Yeast strains were grown at 30°C to an OD600 of 0.8-1.0 in a volume of 2 L of growth medium (YPD/SDC) using a 5 L Erlenmeyer flask with breakers. Cultures were then harvested by centrifugation at 3000g for 5 minutes at 4°C, washed with pre-chilled milli-Q water once and resuspended in lysis buffer (100 mM NaCl, 50 mM Tris-HCl (pH 7.4), 5 mM MgCl2, 0.1% TritonX-100, 10% glycerol, 1 mM DTT and protease inhibitor cocktail (Roche, Germany)), then finally frozen in liquid nitrogen. For 100 OD of cells, 200 μl of lysis buffer was used. For lysis, frozen cell pellets were transferred to a ball mill and beaten for 2-3 minutes using the mixer mill MM 400 (Retsch, Germany). The lysate was then thawed and centrifuged at 20,000g for 30 minutes at 4°C to remove the cell debris. The supernatant was incubated with IgG-SepharoseTM (GE healthcare) for 1 hour at 4°C on rotating wheel and then washed with washing buffer (100 mM NaCl, 50 mM TrisHCl (pH 7.4), 5 mM MgCl2, 0.1% (v/v) NP-40, 10% glycerol, 1 mM DTT). Further, TEV cleavage was performed by adding 2 μl of AcTEV™ protease (Invitrogen) in 1 ml of washing buffer with IgG beads and kept at 16°C for 2 hours on a rotating wheel. Resulting TEV eluate was further incubated with 50 μl anti-FLAG M2 affinity resins (Sigma-Aldrich) for 1 hour at 4°C and washed thrice. Finally, elution was carried out by incubating the washed resins with 600 μl of elution buffer (1X flag peptide in washing buffer) for 45 minutes at 4°C and purified proteins were TCA precipitated prior to SDS-PAGE.

### Mass spec analysis of protein composition by SILAC

For the protein quantification of pre-ribosomes purified by Has1, the stable isotope labeling with amino acids in cell culture (SILAC) method technique was employed as described earlier [Ong et al. 2002]. Briefly, Has1 WT FTpA strain was grown in a “heavy” synthetic complete media containing ^13^C_6_, ^15^N_4_-L-Arginine (Arg-10) and ^13^C_6_, ^15^N_2_-L-Lysine (Lys-8) (Silantes, Munich). The mutants were grown in “light” media with standard amino acids. For standard SILAC experiments, equal amounts of WT and mutant cultures were mixed and subjected to TAP. FLAG elutes were then TCA precipitated, run shortly into 4-12% gradient SDS-PAGE, trypsin digested and peptide masses were analyzed by nLC-MS/MS (in house core facility or at Fingerprints Proteomics Facility, University of Dundee, Scotland). The obtained raw data was processed using MaxQuant software (43). The mass spectrometry proteomics data have been deposited to the ProteomeXchange Consortium via the PRIDE(44) partner repository with the dataset identifier PXD013263.

### Sucrose density gradient analysis of affinity purifications

The eluate from the TAP was loaded onto a 10-40% sucrose density gradient in lysis buffer (100 mM NaCl, 50 mM TrisHCl (pH 7.4), 5 mM MgCl2, 0.1% TritonX-100) and centrifuged at 23 k rpm for 16 hours at 4°C (Beckmann Coulter OptimaTM L-90K ultracentrifuge, SW40 rotor). After the spin, fractions were collected manually and used for protein and RNA analysis.

### Western Blotting

Proteins were isolated from the affinity purification and sucrose gradient fractions of proteins using the TCA precipitation method. And then resolved in 8% SDS-PAGE gel and transferred to Immobilon-FL (Millipore) membrane using wet electro transfer system (Bio-Rad). The membranes were treated with the primary antibodies anti-HA (Abcam), anti-flag, anti-GST (Santo-cruz), anti-Arc1 (AG Hurt) followed by anti-rabbit IgG antibody coupled with Alexa flour 680 (Molecular probes) and scanned on the Odyssay Clx imager (Licor).

### UV Cross-linking and high throughput analysis of cDNAs (CRAC)

CRAC was performed in accordance with the previously described method (21). Briefly, the culture from the strain expressing Has1-HTP was UV cross-linked *in vivo* using megatron UV light for 3 minutes. Following harvesting, the cell pellets were resuspended with lysis buffer (100 mM NaCl, 50 mM Tris-HCl (pH 7.4), 5 mM MgCl2, 0.1% (v/v) NP-40, 10% glycerol, 5 mM β-mercaptoethanol) and lysed by cryo grinding method using the mixer mill MM 400 (Retsch, Germany). The lysate was then subjected to first round of affinity purification with IgG-SepharoseTM (GE healthcare, Germany) for 1 hour at 4°C on the rotating wheel, washed with high salt buffers (1000 mM NaCl, 50 mM Tris-HCl (pH 7.4), 5 mM MgCl2, 0.1% (v/v) NP-40, 10% glycerol and 5 mM β-mercaptoethanol) and then TEV cleaved for 2 hours at 16°C using AcTEV™ protease (Invitrogen, Germany). TEV elute was digested with RNace-It™ Ribonuclease Cocktail (Agilent Technologies, Germany) to remove the non-crosslinked RNAs and purified again with Ni-NTA agarose (Qiagen, Germany) under stringent denaturing conditions using 6 M Guanidium hydrochloride overnight at 4 °C. The beads were then dephosphorylated using the Calf Intestinal Alkaline Phosphatase (CIAP) (Promega, Germany), crosslinked RNAs were sequentially ligated to L3 and bar-coded L5 linkers (Integrated DNA technologies), end-labeled and then eluted using Imidazole (200 mM). Eluates were then resolved on 4-12% gradient SDS-PAGE gel and the band of interest was treated with proteinase-K for 2 hours at 55°C, followed by RNA isolation and RT-PCR. The purified amplicon was subjected to Illumina sequencing (Deep sequencing central facility, Bioquant, Heidelberg) and the sequences obtained were aligned to the yeast genome using Geneious software (www.geneious.com) and Py-CRAC suite (21).

## ACKNOWLEDGEMENTS

We thank Ed Hurt for plasmids and antibodies and his lab members for their helpful discussions. We are grateful to the Dougie Lamont and colleagues from the FingerPrints Proteomics and Mass Spectrometry Facility in Dundee for their help and reliable mass spectrometry service. We also thank the mass spectrometry facility Biochemistry Center (BZH). Many thanks to David Ibberson and the CellNetworks – Deep Sequencing Core Facility, Bioquant Heidelberg for their service. The group of M.K. was funded by the Cluster of Excellence Cell Networks (grant number EXC 81).

## AUTHOR CONTRIBUTIONS

S.G. performed experiments and wrote the manuscript. M.K. designed the study, analyzed data and wrote the manuscript.

