## Supplementary figures for "Two molecules of Has1 RNA helicase function simultaneously in the biogenesis of small and large ribosomal subunits"

**Supplementary figure 1**

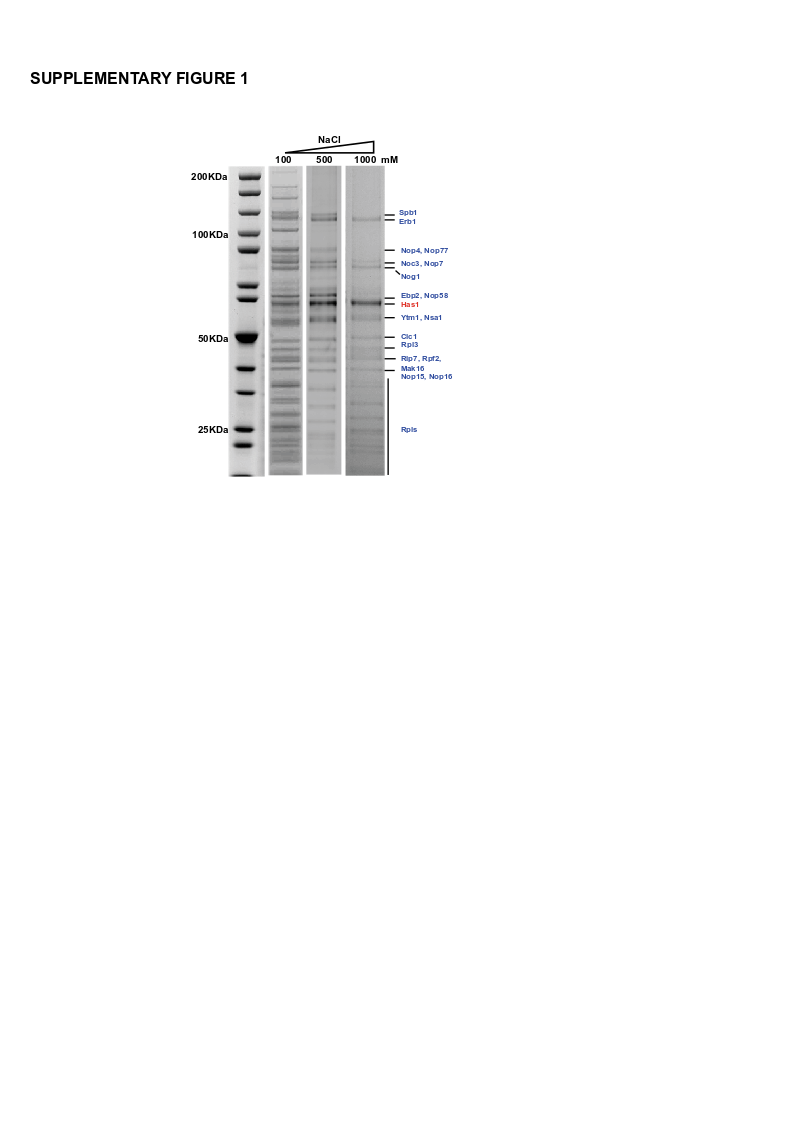

**Supplementary Figure 1.** Protein composition of the Has1 tandem affinity purification after high salt washing. Has1 affinity purification was carried out with high salt washes before elution with the FLAG peptide. The FLAG eluates were resolved in 4-12% gradient SDS-PAGE and stained with Colloidal Coomassie blue. Proteins retained after high salt washing were identified by mass spectrometry.

**Supplementary figure 2**

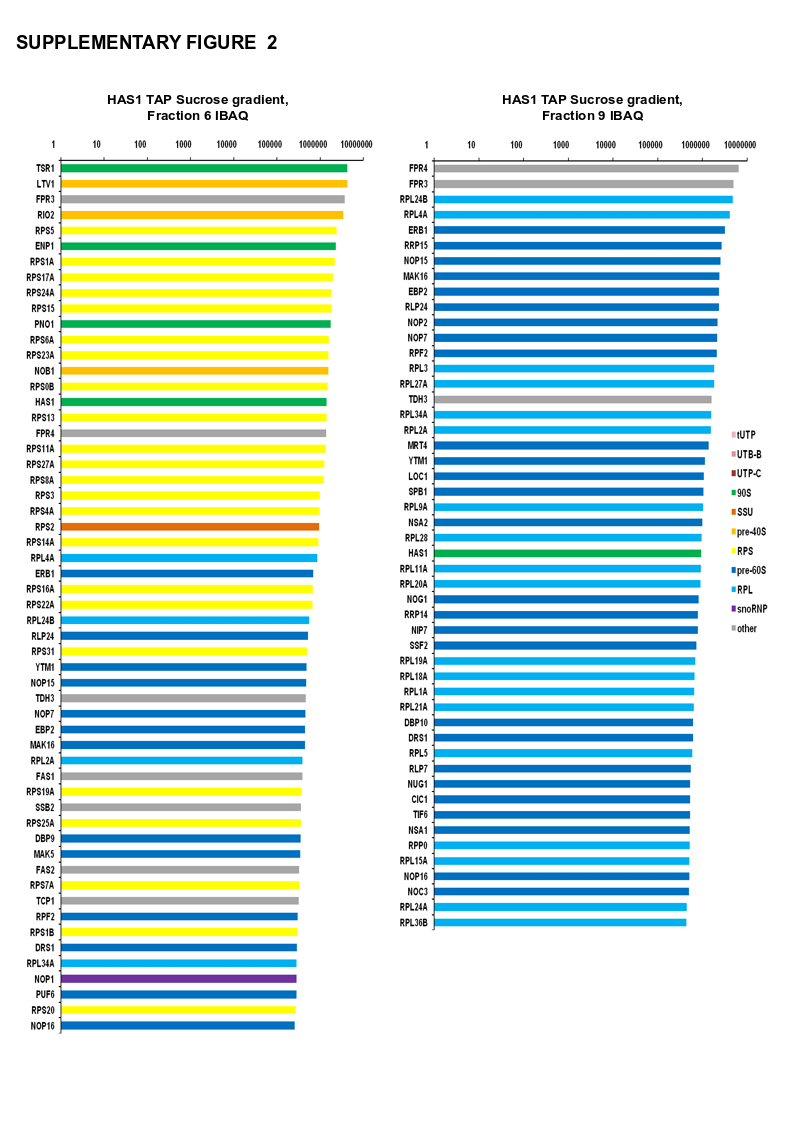

**Supplementary Figure 2.** Graph showing the most abundant proteins (iBAQ values) in the sucrose gradient fractions 6 and 9. Fraction 6 was mostly enriched with pre-40S factors and RPS proteins, while fraction 9 was enriched with pre-60S factors and RPL proteins.

**Supplementary figure 3**

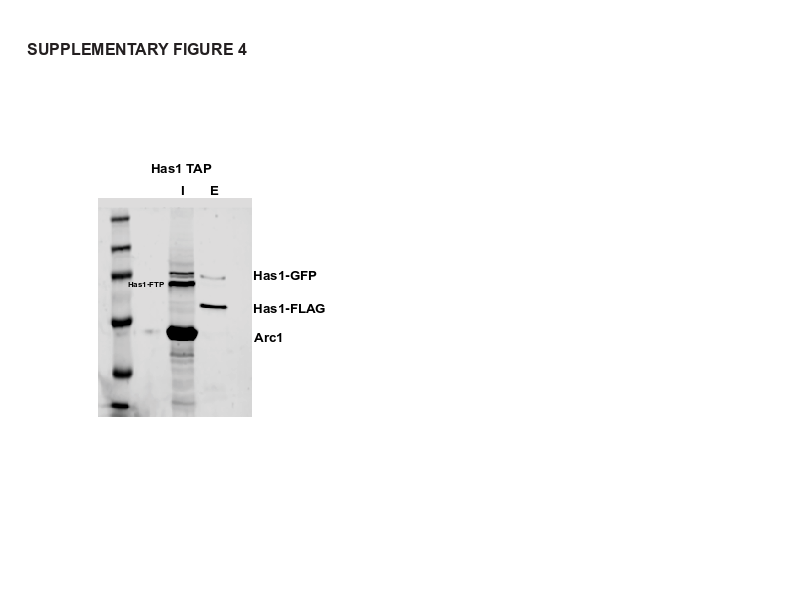

**Supplementary Figure 3.** Yeast strain harboring Has1- GFP and Has1-FTP constructs was subjected to tandem affinity purification and the FLAG eluate was then analyzed by Western blotting using the anti-GFP, anti-FLAG and anti-Arc1 antibodies as indicated. Lane I – input, lane E – FLAG eluate.
