## Supplementary tables 3-7 for "Two molecules of Has1 RNA helicase function simultaneously in the biogenesis of small and large ribosomal subunits"

**Supplementary Table 3: Has1 WT binding sites in rRNA**

| **Peak** | **RDNA37-1 (nts)** | **Helix number** | **Other factors binding in the helix** |
| --- | --- | --- | --- |
| 1 | 1908-1966 | 31,33 | ENP1, RIO2 |
| 2 | 2012-2058 | 37, 38, 39 | LTV1 |
| 3 | 2119-2211 | 32, 40, 41 | NOB1, LTV1 |
| 4 | 2939-2992 | 7,5,8,9 | NOP12 |
| 5 | 3492-3532 | 17 16 21 | ERB1 |
| 6 | 5791-5806 | 79 | - |
| 7 | 6555-6633 | 98 101 | - |

**Supplementary Table 4: Has1 DAAD binding sites in rRNA**

| **Peak** | **RDNA37-1 (nts)** | **Helix number** | **Other factors binding in the helix** |
| --- | --- | --- | --- |
| 1 | 1227-1273 | 17, 18 | - |
| 2 | 3533-3572 | 21 22 | ERB1 |
| 3 | 5397-5470 | 65 66 | - |
| 4 | 5765-5839 | 79 | - |
| 5 | 6114-6136 | 89 | - |
| 6 | 6555-6608 | 98 101 | - |

**Supplementary Table 5: Plasmid constructs used in this study.**

| **Plasmid** | **Description** | **Reference** |
| --- | --- | --- |
| pRS425 | 2μ, Episomal, LEU2 | Christianson *et al.,* 1992 |
| pMK071 | pFA6a-GFP(S265T)::NatNT2 | Bassler *et al.,* 2010 |
| pMK424 | pHTP::HIS3MX6 | This study |
| pMK604 | pRS425-URA3-LEU2d-18S(604nt)-MS2bs-Bo-2 enh-short prom | Kos-Braun et al., |
| pMK605 | pRS425-URA3-LEU2d-18S(1142nt)-MS2bs-Bo-2 enh-short prom | Kos-Braun et al., |
| pMK606 | pRS425-URA3-LEU2d-18S(1780nt)-MS2bs-Bo-2 enh-short prom | Kos-Braun et al., |
| pMK607 | pRS425-URA3-LEU2d-C2-MS2bs-Bo-2 enh-short prom | Kos-Braun et al., |
| pMK608 | pRS425-URA3-LEU2d-25S(421nt)-MS2bs-Bo-2 enh-short prom | Kos-Braun et al., |
| pMK609 | pRS425-URA3-LEU2d-25S(1453nt)-MS2bs-Bo-2 enh-short prom | Kos-Braun et al., |
| pMK610 | pRS425-URA3-LEU2d-25S(2362nt)-MS2bs-Bo-2 enh-short prom | Kos-Braun et al., |
| pMK611 | pRS425-URA3-LEU2d-35S-MS2bs-Bo-2 enh-short prom | Kos-Braun et al., |
| pMK621 | pFA6a-FTP::NatNT2 | Gift from Ed Hurt lab |
| pMK623 | pRS415-LEU2-Has1WT-FTP | This study |
| pMK632 | pMK140-TetO7-Ubi-Leu-3HA::NatNT2 | This study |
| pMK633 | pMK140-TetO7-Ubi-Tyr-3HA::NatNT2 | This study |
| pMK634 | pMK140-TetO7-Ubi-Ile-3HA::NatNT2 | This study |
| pMK635 | pMK140-TetO7-Ubi-Ala-3HA::NatNT2 | This study |
| pMK774 | pFA6a-FTP::HIS3MX4 | Gift from Ed Hurt lab |
| pMK813 | pRS425-URA3-LEU2d-post Dsite-25S(421nt)-MS2bs-B0-2 enh | This study |
| pMK814 | pRS425-URA3-LEU2d-A3-25S(421nt)-MS2bs-B0-2 enh | This study |
| pMK815 | pRS425-URA3-LEU2d-5.8S-25S(421nt)-MS2bs-B0-2 enh | This study |
| pMK816 | pRS425-URA3-LEU2d-E site-25S(421nt)-MS2bs-B0-2 enh | This study |
| pMK830 | pRS425-URA3-LEU2d-5.8S(155nt)-MS2bs-B0-2 enh-18Stag | This study |
| pMK831 | pRS425-URA3-LEU2d-C2-MS2bs-B0-2 enh-18Stag | This study |
| pMK832 | pRS425-URA3-LEU2d-25S (421nt)-MS2bs-B0-2 enh-18Stag | This study |
| pMK866 | pRS415-LEU2-Has1KA-FTP | This study |
| pMK867 | pRS415-LEU2-Has1DAAD-FTP | This study |
| pMK868 | pRS415-LEU2-Has1AAA-FTP | This study |
| pMK870 | pRS425-URA3-LEU2d-18S-postA2-MS2bs-B0-2enh-18S tag | This study |
| pMK869 | pRS425-URA3-LEU2d-18S-beforeA2-MS2bs-B0-2enh-18S tag | This study |
| pMK140 | pMK140-TetO7-3HA::NatNT2 | This study |
| pMK014 | pFA6a-GST::HIS3MX6 | Longtine *et al.,* 1998 |
| pMK711 | pRS315-Has1-eGFP | This study |
| pMK560 | pRS415-LEU2-FTP | This study |

**Supplementary Table 6: Yeast strains used in this study.**

| **YMK number** | **Genotype** | **Plasmid used** | **Reference** |
| --- | --- | --- | --- |
| YMK118 | CEN-PK2-1C,MATa; HIS3 ∆1; LEU2-3,112; TRP1-289; URA3-52; MAL2-8C; SUC2, LYS2:: tTA, URA3::PCMVtetR'-SSN6 URA3-K.l. | - | Alexander et al., 2010 |
| YMK444 | CEN-PK2-1C,MATa; PtetO7-Ubi-Leu-3HA-Has1-NatNT2 | pMK632 | Gnanasundram et al., 2015 |
| YMK604 | CEN-PK2-1C,MATa;Has1-FTP:: HIS3MX6 | pMK774 | This study |
| YMK539 | CEN-PK2-1C,MATa; Arg4∆, PtetO7-Ubi-Leu-3HA-Has1-NatNT2 | - | This study |
| YMK592 | CEN-PK2-1C,MATa; Arg4∆, PtetO7-Ubi-Leu-3HA-Has1-NatNT2+ pRS415-Has1 WT-FTP | pMK623 | This study |
| YMK593 | CEN-PK2-1C,MATa; Arg4∆, PtetO7-Ubi-Leu-3HA-Has1-NatNT2+ pRS415-Has1 KA-FTP | pMK866 | This study |
| YMK594 | CEN-PK2-1C,MATa; Arg4∆, PtetO7-Ubi-Leu-3HA-Has1-NatNT2+ pRS415-Has1 DAAD-FTP | pMK867 | This study |
| YMK595 | CEN-PK2-1C,MATa; Arg4∆, PtetO7-Ubi-Leu-3HA-Has1-NatNT2+ pRS415-Has1 AAA-FTP | pMK868 | This study |
| YMK146 | CEN-PK2-1C,MATa; URA3- replaced with Hygromycin | - | This study |
| YMK688 | CEN-PK2-1C,MATa; URA3- replaced with Hygromycin, Has1-FTP::NatNT2 | pMK621 | This study |
| YMK693 | CEN-PK2-1C,MATa; URA3- replaced with Hygromycin, Has1-FTP::NatNT2 + pMK603-A0- MS2bs-B0-2enh) | pMK603 | This study |
| YMK694 | CEN-PK2-1C,MATa; URA3- replaced with Hygromycin, Has1-FTP::NatNT2 + pMK604 (18S [604nt]- MS2bs-B0-2enh) | pMK604 | This study |
| YMK695 | CEN-PK2-1C,MATa; URA3- replaced with Hygromycin, Has1-FTP::NatNT2 + pMK605 (18S [1142nt]- MS2bs-B0-2enh) | pMK605 | This study |
| YMK696 | CEN-PK2-1C,MATa; URA3- replaced with Hygromycin, Has1-FTP::NatNT2 + pMK606- (18S [1780nt]- MS2bs-B0-2enh) | pMK606 | This study |
| YMK697 | CEN-PK2-1C,MATa; URA3- replaced with Hygromycin, Has1-FTP::NatNT2 + pMK607-(C2- MS2bs-B0-2enh) | pMK607 | This study |
| YMK698 | CEN-PK2-1C,MATa; URA3- replaced with Hygromycin, Has1-FTP::NatNT2 + pMK608 (25S [421nt]- MS2bs-B0-2enh) | pMK608 | This study |
| YMK699 | CEN-PK2-1C,MATa; URA3- replaced with Hygromycin, Has1-FTP::NatNT2 + pMK609 (25S [1453nt]- MS2bs-B0-2enh) | pMK609 | This study |
| YMK700 | CEN-PK2-1C,MATa; URA3- replaced with Hygromycin, Has1-FTP::NatNT2 + pMK610 (25S [2362nt]- MS2bs-B0-2enh) | pMK610 | This study |
| YMK701 | CEN-PK2-1C,MATa; URA3- replaced with Hygromycin, Has1-FTP::NatNT2 + pMK611 (35S-MS2bs-B0-2enh) | pMK611 | This study |
| YMK878 | CEN-PK2-1C,MATa; URA3- replaced with Hygromycin, Has1-FTP::NatNT2 + pMK830- 5.8S[155nt]-MS2bs-Bo-Rnt1-18Stag | pMK830 | This study |
| YMK879 | CEN-PK2-1C,MATa; URA3- replaced with Hygromycin, Has1-FTP::NatNT2 + pMK831- C2-MS2bs-Bo-Rnt1-18Stag | pMK831 | This study |
| YMK880 | CEN-PK2-1C,MATa; URA3- replaced with Hygromycin, Has1-FTP::NatNT2 + pMK832- 25S[421nt]-MS2bs-Bo-Rnt1-18Stag | pMK832 | This study |
| YMK881 | CEN-PK2-1C,MATa; URA3- replaced with Hygromycin, Has1-FTP::NatNT2 + pMK813- post D site-25S[421nt]-MS2bs-B0-2enh | pMK813 | This study |
| YMK882 | CEN-PK2-1C,MATa; URA3- replaced with Hygromycin, Has1-FTP::NatNT2 + pMK814- A3 site-25S[421nt]-MS2bs-B0-2enh | pMK814 | This study |
| YMK883 | CEN-PK2-1C,MATa; URA3- replaced with Hygromycin, Has1-FTP::NatNT2 + pMK815- 5.8S-25S[421nt]-MS2bs-B0-2enh | pMK815 | This study |
| YMK884 | CEN-PK2-1C,MATa; URA3- replaced with Hygromycin, Has1-FTP::NatNT2 + pMK816- E site-25S[421nt]-MS2bs-B0-2enh | pMK816 | This study |
| YMK646 | CEN-PK2-1C,MATa; Has1-HTP::HIS3MX6 | pMK424 | This study |
| YMK673 | CEN-PK2-1C,MATa; + pRS415-Has1-DAAD-HTP-LEU2 |  | This study |
| YMK845 | CEN-PK2-1C,MATa; Has1-eGFP::NatNT2 | pMK071 | This study |
| YMK261 | CEN-PK2-1C,MATa ; tetO7::3HA-Has1::NatNT2 | pMK140 | This study |
| YMK955 | CEN-PK2-1C,MATa; Has1-GST::HIS3MX6 | pMK014 | This study |
| YMK949 | CEN-PK2-1C,MATa; URA3- replaced with Hygromycin, Has1-FTP::HIS3MX6 | - | This study |

**Supplementary table 7: Oligonucleotides used for Northern hybridization.**

| Probe used for detecting | Sequence |
| --- | --- |
| 35S | GCTGCTCACCAATGGAATC |
| 27SA2-A3 | GCAAAGATATGAAAACTCCAC |
| 27S A3-B1 | GTTCCAGTTACGAAAATTCTTGT |
| 27C1-C2 | GTTCGCCTAGACGCTCTCTT |
| 20S | CGGTTTTAATTGTCCTA |
| 25S | CTCCGCTTATTGATATGC |
| 18S | CATGGTTAATCTTTGAGAC |
| 7S | GGCCAGCAATTTCAAGTTA |
| 5.8S | GCGTTCTTCATCGATGC |
| MS2 | GTCTTTCTATCGACATGGGTG |
| SCR1 | ATCCCGGCCGCCTCCATCAC |
| 18S tag | GAGGATCCAGGTTTGTC |
